# RT-QuIC amplification of CWD prions in winter ticks (*Dermacentor albipictus*) collected from North American elk (*Cervus canadensis*) in a CWD-endemic area

**DOI:** 10.1101/2021.06.04.447178

**Authors:** NJ Haley, DM Henderson, K Senior, M Miller, R Donner

## Abstract

Chronic wasting disease (CWD) is a progressive and fatal spongiform encephalopathy of deer and elk species, caused by a misfolded variant of the normal prion protein. Horizontal transmission of the misfolded CWD prion between animals is thought to occur through shedding in saliva and other forms of excreta. The role of blood in CWD transmission is less clear, though infectivity has been demonstrated in various blood fractions. Blood-feeding insects, including ticks, are known vectors for a range of bacterial and viral infections in animals and humans, though to date there has been no evidence for their involvement in prion disease transmission. In the present study, we evaluated winter ticks (*Dermacentor albipictus*) collected from 136 North American elk (*Cervus canadensis*) in a CWD-endemic area for evidence of CWD prion amplification using the real time quaking-induced conversion assay (RT-QuIC). Although 30 elk were found to be CWD-positive (22%) postmortem, amplifiable prions were found in just a single tick collected from an elk in advanced stages of CWD infection, with some evidence for prions in ticks collected from elk in mid-stage infection. These findings suggest that further investigation of ticks as reservoirs for prion disease may be warranted.

**Importance:** This study reports the first finding of detectable levels of prions linked to chronic wasting disease in a tick collected from a clinically infected elk. Using the real time quaking-induced conversion assay (RT-QuIC), evidence of amplifiable CWD prions was also found in ticks collected from elk in earlier stages of disease. Observed levels were at the lower end of our detection limits, though our findings suggest that additional research evaluating ticks collected from animals in late-stage disease may be warranted to further evaluate the role of ticks as potential vectors of chronic wasting disease.

## Introduction

Chronic wasting disease (CWD) is a progressive and ultimately fatal neurodegenerative disease in cervids, including whitetail deer (*Odocoileus virginianus*), mule deer (*Odocoileus hemionus*), and North American elk (wapiti, *Cervus canadensis*) (1, 2). Like other diseases in this category – the transmissible spongiform encephalopathies or TSEs – CWD is caused by an infectious, misfolded variant of the normal cellular prion protein (PrP^C^) that is often designated as PrP^CWD^ (3, 4). First identified in captive mule deer in northern Colorado and southern Wyoming in the late 1960s, CWD has now been reported in farmed and/or free ranging cervids in 26 U.S. states, three Canadian provinces, South Korea, Norway, Sweden, and Finland (5–9).

*In vivo* studies, using both deer and mouse bioassay, and *in vitro* amplification assays have identified infectious prions or PrP^CWD^ in various bodily fluids and other forms of excreta, including saliva, blood, urine, and feces (10–19). In nature, it is thought that saliva is an important means of direct animal to animal transmission, while urine, feces, and decomposing carcasses may all play important roles in environmental contamination and subsequent exposure and transmission (2, 20, 21). The role of blood in the transmission of CWD and other prion diseases is less clear, though various blood fractions have been found to convey prion infection both in experimental studies in animal models and through rare natural infections in humans (11, 22–24). Importantly, the role of blood feeding insects, including ticks, as vectors for prion transmission in nature has not been extensively evaluated.

Winter ticks (*Dermacentor albipictus*) are common external parasites found on moose, caribou, elk, and other large herbivores, and are widely distributed across North America, Mexico, and Central America (25). They may be found in very high densities on moose, sometimes numbering in the tens of thousands on a single animal, and have been linked to declining moose populations across North America (26). Winter ticks are considered a “one-host” tick, taking several blood meals from a single host while progressing through larval and nymph stages to adulthood, though they are likely to seek a new host if dislodged prematurely from their primary host (27, 28). As a single-host tick, they have only rarely been implicated as vectors of disease, notably hemoparasites such as *Anaplasma marginale* and *Babesia duncani* (29, 30). Paradoxically, this one-host lifecycle makes them ideal for evaluating tick species as potential reservoirs for CWD and other prions, as every blood meal over the course of their development is putatively taken from a single host, thus amplifying their exposure to the agent.

As part of a prior study evaluating the practicality of antemortem rectal biopsy testing in managing CWD in ranched elk (31), we collected adult winter ticks from elk in an area with a high prevalence of CWD. Of the 136 elk sampled, 30 were found to be CWD-positive either antemortem or postmortem (22%), at various stages of disease based on the diverse appearance of PrP^CWD^ in diagnostic tissues and apparent clinical symptoms suggestive of late-stage infection. In the present study, we blindly evaluated these ticks using the real-time quaking induced conversion assay (RT-QuIC), an assay used widely to amplify PrP^CWD^ in a range of substrates, including rectal biopsies in the parent study (14–17, 31–36). We hypothesized that amplifiable PrP^CWD^ would be detected in ticks collected from CWD-positive elk, with ticks from elk in more advanced stages of disease having higher rates of amplification. We found PrP^CWD^ amplification in a single tick collected from an elk in an advanced stage of CWD, with evidence for amplifiable prions in ticks from elk in mid-stage CWD. Though the biological relevance of our findings remains unknown, they suggest that winter ticks, and potentially other tick species, could serve as low-level reservoirs for CWD in nature.

## Materials and Methods

### Ethics statement

The animals from which ticks were collected in this study were handled humanely in accordance with Midwestern University’s Animal Care and Use Committee, approval #2814.

### Elk study population

The elk involved in the study were part of a private, closed herd living on 3500 acres of fenced-in land in northwestern Colorado. A thorough description of the study population and area may be found elsewhere (32). Winter ticks were collected from 136 bull and cow elk ranging in age from calves to 14 years. A mixture of 132M/L genotypes were represented among the elk sampled. Postmortem tissues including both retropharyngeal lymph node (RLN) and brainstem at the level of the obex were evaluated for evidence of CWD infection using immunohistochemistry as described previously (31).

### Tick collection and processing

Adult winter ticks were collected cleanly, using single-use gloves, from the pinnae of elk and stored at −80 °C for approximately two years until processing. At that time, a single tick was recovered from storage, weighed, homogenized in phosphate-buffered saline (PBS) at 1% w/v using a bead homogenizer and frozen prior to analysis. After thawing, homogenates were further diluted for experimental evaluation as described in the following section. Ticks collected from three elk calves with the 132LL genotype were identified to serve as putative negative controls throughout the study.

### Optimization of the real time quaking-induced conversion assay (RT-QuIC) for tick substrate

To find a roughly optimal tick starting dilution without sacrificing dilutional sensitivity, a pilot experiment was conducted to assess the amplification ability of PrP^CWD^ from known a CWD-positive RLN in three dilutions of tick substrate: 1%, 0.1%, and 0.01% (10^−2-4^) tick homogenates prepared in PBS, compared to RLN homogenized in PBS alone. First, RLN from a known CWD-positive whitetail deer was homogenized in PBS at a 10% (10^−1^) w/v dilution. Aliquots of this RLN preparation were then diluted serially ten-fold from 10^−4^ to 10^−7^ in either PBS or 10^−2^, 10^−3^, or 10^−4^ homogenates of ticks collected from 132LL genotype elk calves. Amplification was performed using a truncated form of recombinant Syrian hamster PrP (SHrPrP, residues 90-231) as a conversion substrate. The recombinant SHrPrP was prepared off-site, frozen at −80 °C and thawed slowly just prior to use in each experiment. Two μl of each of the preparations was added to 98 μl of RT-QuIC master mix (50 mM NaPO4, 350 mM NaCl, 1.0 mM EDTA, 10mM thioflavin T [ThT], and 0.1 mg/ml SHrPrP). Individual sample dilutions were repeated in triplicate, in two separate experiments, in a 96-well, optical-bottom plate, sealed and incubated in a BMG Labtech PolarstarTM fluorimeter at 42°C for 48 hours (192 cycles, 15 minutes each) with intermittent shaking. Cycle parameters included 1-minute shakes (700 rpm, double orbital pattern) interrupted by 1 minute rest periods, with ThT fluorescence measurements (450 nm excitation and 480 nm emission) taken every 15 minutes with the gain set at 1800. The relative fluorescence units (RFU) for each triplicate sample were progressively monitored against time with orbital averaging and 20 flashes/well at the 4 mm setting. Time to amplification was determined based on individual replicates crossing an experimental threshold calculated as ten standard deviations above the mean fluorescence of all sample wells across amplification cycles 2-8, as described in previous studies (31, 32, 37).

### RT-QuIC evaluation of ticks collected from elk

After identifying appropriate dilution levels for tick homogenates and RLN positive controls, study ticks were individually homogenized and evaluated by RT-QuIC using parameters described above. Study plates included (1) positive RLN controls repeated in triplicate, consisting of a 10^−5^ dilution of RLN in a 10^−3^ tick homogenate in PBS – which typically amplified between cycle number 80-100, (2) individual tick samples prepared at a10^−3^ dilution evaluated in triplicate, (3) three unique negative controls, each prepared in triplicate (9 total replicates), consisting of tick homogenates from three elk calves with 132LL genotypes, also prepared at a 10^−3^ dilution, and (4) unspiked SHrPrP, also in triplicate. Tick homogenates were analyzed blindly, without information on source animal CWD status during tick processing, amplification, and data analysis stages, with CWD status revealed only upon completion of the analysis. All samples were repeated in triplicate in two separate experiments for a total of six replicates.

Criteria for identification of positive samples was determined *a priori* and was consistent with previous studies in our laboratories (31, 32, 37). A replicate well was considered positive when the relative fluorescence crossed a pre-defined threshold as described above. Positive samples were those which crossed the threshold in ≥3 out of 6 replicates; samples with 1-2 of 6 replicates positive were considered “suspects.” Plates were disqualified if any of the positive control replicates failed to amplify, or if amplification was observed in any of the various negative control replicates.

Amplification rates of positive and suspect samples were subsequently compared to a standard curve generated in the preliminary experiment, using RLN serially diluted into a 10^−3^ dilution of tick homogenate.

### Statistical analysis

In our preliminary experiment, differences between amplification rates assessing RLN dilutions in various tick backgrounds were compared using two-tailed Student’s t-tests. Approximate RLN dilutional equivalents of RT-QuIC positive and RT-QuIC suspect ticks were calculated by comparing the mean amplification rate of the six replicates to a non-linear fit of data from RLN diluted in 10^−3^ tick homogenate in preliminary tick dilution experiments, developed using GraphPad Prism 8.4.1 software. Comparison of the likelihood of “suspect” amplification occurring in ticks collected from CWD-negative and CWD-positive elk was performed using a two-tailed Fisher exact test.

## Results

### Summary of Source Elk Testing

A total of 30 of the 136 elk providing ticks for the present study were identified as CWD-positive via postmortem immunohistochemistry of RLN and obex tissue (22%). Based on the distribution of PrP^CWD^ in these tissues as well as clinical presentation, infected animals were estimated to be in various stages of disease from early and pre-clinical to late stage and symptomatic. A breakdown of estimated disease stages can be found in **Table 1**.

**Table 1:** Summary of elk providing tick samples for the present study. Genotype is based on the amino acid coded for at position 132 of the elk prion gene, Prnp. Animals with a 132MM genotype are comparatively more susceptible, while those with a 132ML or 132LL genotype are less likely to be found CWD positive in natural and experimental conditions. Disease stage estimates are based on immunohistochemical detection of PrP^CWD^ in the elk’s brain (B) and/or retropharyngeal lymph node (RLN), and clinical signs consistent with late-stage infection, including behavior abnormalities and poor body condition scores.

| CWD Status | Total | Mean Age (Range) | Prnp Genotype |  |  | Estimated Disease Stage |  |  |
| --- | --- | --- | --- | --- | --- | --- | --- | --- |
|  |  |  |  |  |  | Early | Middle | Late |
|  |  |  | 132MM | 132ML | 132LL | B or RLN (+) | B and RLN (+) | B and RLN (+) and Symptomatic |
| Positive | 30 | 5.2 (2 – 14) | 18 | 10 | 2 | 5 | 24 | 1 |
| Negative | 106 | 3.07 (1 – 13) | 41 | 49 | 16 |  |  |  |

### Optimized dilution of ticks for use in RT-QuIC

Pilot experiments found that RLN homogenates prepared in both 10^−3^ and 10^−4^ dilutions of tick homogenate amplified at rates most similar to RLN homogenates prepared in PBS alone. Tick homogenates of 10^−3^ were selected for primary studies based on this similarity and to avoid sacrificing potential dilutional sensitivity in 10^−4^ dilution of tick homogenates. Retropharyngeal lymph node dilutions down to 10^−7^ amplified in a 48hr assay, representing the lower limits of detection for our study. (**Figure 1**)

**Figure 1:**
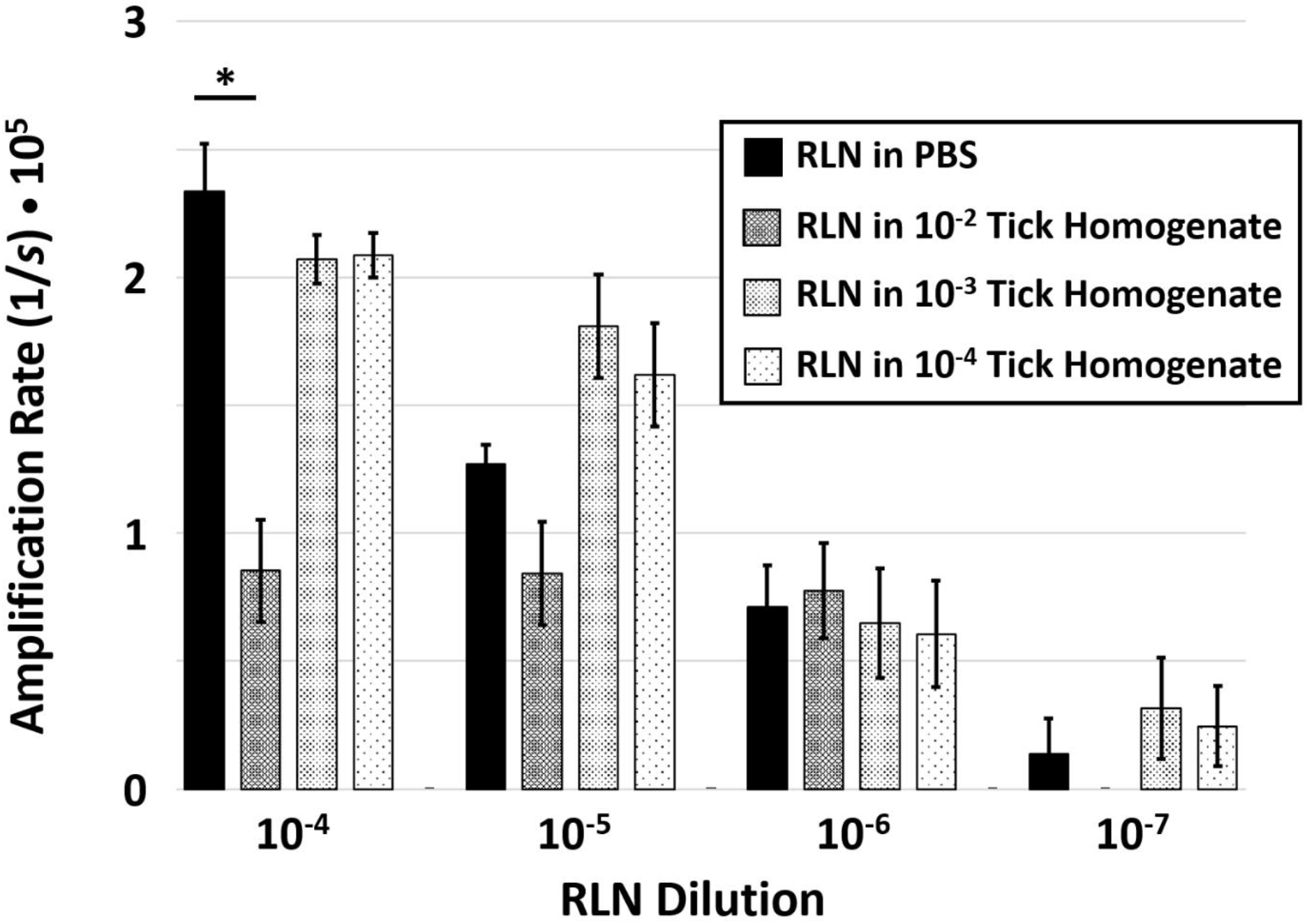
Amplification rates of CWD-positive retropharyngeal lymph node (RLN) dilutions in phosphate-buffered saline (PBS) or tick homogenate dilutions. Amplification rates were calculated as the average inverse of the time, in seconds, to amplification threshold in six replicate wells across two experimental plates. Standard error bars are shown. Using a two-tailed Student’s t-test, a significant reduction in amplification (*, p ≤ 0.01) was observed in the 10^−4^ dilution of RLN in PBS compared to RLN diluted in 10^−2^ tick homogenate.

### RT-QuIC amplification of CWD prions in ticks

In the primary experiments, results from four amplification plates were discarded due to amplification in a negative control replicate; samples from these plates were subsequently repeated with no amplification among negative controls.

Among the 136 ticks analyzed from elk, a single positive tick was identified based on amplification in 3/6 replicates across two blinded experiments. This tick was collected from a CWD-positive elk in late-stage symptomatic disease, and the mean amplification rate was approximately equal to a 10^−6.5^ dilution of CWD-positive RLN (**Figure 2**). Thirteen ticks demonstrated amplification considered as “suspect,” with 1-2 of 6 replicates showing amplification, all with mean amplification rates between 10^−6.9^ and 10^−7.2^ RLN dilutional equivalents (e.g., approximating or beyond the lower range of our non-linear regression fit). Of these suspects, six were from CWD-positive elk in middle stages of disease, while seven were from CWD-negative elk. Ticks from CWD-positive elk were significantly more likely to be considered “suspect” than CWD-negative elk using a two-tailed Fisher exact test (6/29 vs. 7/106, *P* = 0.0337). No evidence of amplification was observed in ticks from 23/30 CWD-positive elk and 106/113 CWD-negative elk. (**Table 2**)

**Table 2:** Summary of RT-QuIC results from ticks collected from CWD-positive and -negative elk. Disease stage estimates were again based on immunohistochemical detection of PrP^CWD^ in the elk’s brain (B) and/or retropharyngeal lymph node (RLN), and clinical signs consistent with late-stage infection, including behavior abnormalities and poor body condition scores.

| Elk CWD Status | Total | RT-QuIC of Winter Ticks |  |  |
| --- | --- | --- | --- | --- |
|  |  | No amplification | Suspect | Positive |
| Early (B or RLN) | 5 | 5 | 0 | 0 |
| Middle (B and RLN) | 24 | 18 | 6 | 0 |
| Late (B and RLN and Symptomatic) | 1 | 0 | 0 | 1 |
| Negative | 106 | 99 | 7 | 0 |

**Figure 2:**
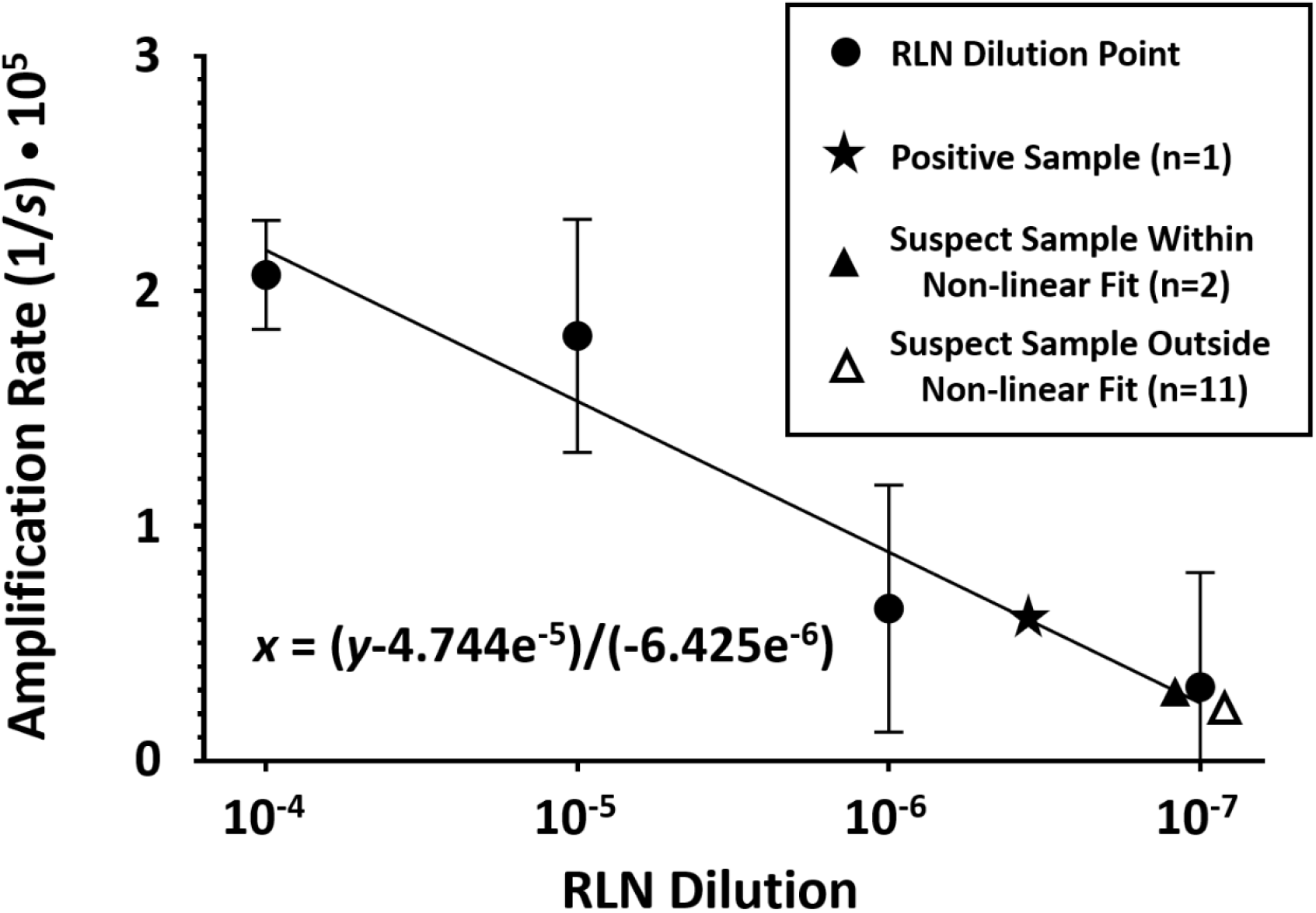
Non-linear regression fit of data from CWD-positive retropharyngeal lymph node (RLN) dilutions in a 10^−3^ tick homogenate. Amplification rates were calculated as the inverse of the average time, in seconds, to the amplification threshold in six replicate wells across two experimental plates. Data points shown include RLN dilutions with standard error bars as well as a single RT-QuIC positive sample and suspect samples both within and outside the range of RLN dilutional data. The slope of the non-linear regression line is indicated as x = (y-4.744^−5^)/(−6.425e^−6^).

## Discussion

With the sharp global decline in reported sheep scrapie and bovine spongiform encephalopathy cases (38, 39), chronic wasting disease of cervids remains one of the most important mammalian prion diseases identified to date – occurring in both free-ranging and farmed populations of deer and related species across North America and now Scandinavia (40). Over the past two decades, much has been learned about the mechanisms of CWD transmission, primarily through the evaluation of blood and other body fluids in multi-year long cervid or murine bioassay experiments (10, 11, 13, 22, 41–46). More recently, *in vitro* amplification assays, which typically offer results in several days, have sought to supplant bioassay as the dominant testing modality for PrP^CWD^ in various forms of excreta (13–17, 47). These assays have been systematically used to evaluate the onset, duration, and severity of PrP^CWD^ shedding in saliva, urine, and feces. The real time quaking-induced conversion assay (RT-QuIC) in particular has been shown to be sensitive, specific, and highly reproducible when evaluating biological samples for CWD prions (31, 32, 37). In the present study, we evaluated ticks collected from CWD-negative elk and elk in various stages of CWD ranging from early pre-clinical infections to late-stage symptomatic disease, to better characterize their potential role as vectors of CWD transmission.

We began by roughly optimizing tick homogenate concentrations for use in the RT-QuIC assay, by comparing the amplification of PrP^CWD^ from known positive RLN tissue in varying dilutions of background tick homogenate. We found that RLN diluted in a 10^−3^ preparation of tick homogenate allowed for amplification of PrP^CWD^ similar to RLN diluted in PBS alone, without potentially sacrificing diluted sensitivity. We went on to blindly evaluate ticks collected from 136 elk, in triplicate and in two independent experiments. We identified a single tick as RT-QuIC positive based on amplification in three of six replicates; this tick had been collected from an elk in terminal stages of CWD based on clinical and pathological findings. Several ticks from both CWD-negative and CWD-positive elk were found to be RT-QuIC “suspects”, based on amplification in one or two of six replicates. Ticks from CWD-positive elk, specifically those in mid-stage CWD, were significantly more likely to be considered RT-QuIC suspects than those from CWD-negative elk. The rates of amplification observed in suspect cases were at the lower limits of detection for the assay, and the biological relevance of true PrP^CWD^ amplified from RT-QuIC suspect and positive ticks is not known. Importantly, we cannot completely rule out dermal or environmental contamination of the ticks, from dust or excreta for example. We likewise cannot rule out that ticks from CWD-negative elk which showed amplification had not recently relocated from a CWD-positive elk, though this seems unlikely. We must therefore be cautious not to weigh these findings too heavily, and instead reinforce the importance of adequate negative controls and proper experimental blinding in future prion amplification studies (48).

Admittedly, additional experiments on *D. albipictus* and other tick species from deer and elk, especially those in later stages of CWD, are necessary to further explore the role of ticks as prion reservoirs. A recent study in a hamster model of CWD found that the Rocky Mountain wood tick, *Dermacentor andersoni*, is unlikely to transmit biologically relevant levels of prions to a naïve host after consuming a blood meal (49). It is important to highlight that, unlike *D. andersoni*, the winter ticks evaluated in the present study are one-host ticks (25). These single host ticks represent ideal targets to assess PrP^CWD^ accumulation, as they had consumed several blood meals prior to collection, yet they are arguably unlikely to transmit infectious agents between hosts in nature. Though they have only rarely been suspected as disease vectors, winter ticks are thought to seek new hosts when feeding is interrupted, for example when removed through grooming or, understandably, following the demise of the host due to e.g., CWD (25, 26, 29, 50). Vertical transmission of adult *D. andersoni* ticks to neonatal moose calves has also been reported (28). For those reasons, a better understanding of the biological relevance of any detectable PrP^CWD^ in this and other species of ticks is warranted.

In summary, we report that RT-QuIC may serve as a useful tool for evaluating the role of ticks and other insects as reservoirs of PrP^CWD^. Amplifiable levels of PrP^CWD^ in the present study were low, and likely limited to ticks collected from animals in later stages of disease. Additional studies focusing on insect vectors feeding on terminally infected cervids and the biological relevance of any detectable CWD prions in these vectors are warranted to more fully characterize the role of external parasites in prion transmission.

## Acknowledgements

This project was funded through a Midwestern University Intramural Grant, in coordination with the Colorado Department of Agriculture, CWDEvolution LLC, and the United States Department of Agriculture Animal and Plant Health Inspection Service. The authors would like to thank the ranch owners, as well as Aaron Lehmkuhl and Bruce Thomsen with the Animal and Plant Health Inspection Service of the USDA, Dan Love, Ed Kline, Keith Roehr, Dave Dice, and Wayne East with the Colorado Department of Agriculture, Jeffrey Christiansen with Colorado State University’s Prion Research Center, and Mike Miller of the Colorado Division of Parks and Wildlife. Without the gracious assistance of these groups and individuals, this project could not have been undertaken.

## Disclosure of Potential Conflicts of Interest

DMH is the owner and operator of CWDEvolution LLC, a private company which produces recombinant prion protein for use in the RT-QuIC assay. DMH had no role in study design or analysis.

## Funding

This work was supported by a Midwestern University Intramural Grant #30-2022-8141.

## References

1. Williams ES, Young S. 1980. Chronic wasting disease of captive mule deer: a spongiform encephalopathy. J Wildl Dis 16:89–98.

2. Haley NJ, Hoover EA. 2015. Chronic Wasting Disease of Cervids: Current Knowledge and Future Perspectives. Annu Rev Anim Biosci doi:10.1146/annurev-animal-022114-111001.

3. Ironside JW, Ritchie DL, Head MW. 2017. Prion diseases. Handb Clin Neurol 145:393–403.

4. Wolfe LL, Spraker TR, Gonzalez L, Dagleish MP, Sirochman TM, Brown JC, Jeffrey M, Miller MW. 2007. PrPCWD in rectal lymphoid tissue of deer (Odocoileus spp.). J Gen Virol 88:2078–82.

5. ProMED-Mail. 2019. CHRONIC WASTING DISEASE - SWEDEN: (NORRBOTTEN) MOOSE, FIRST CASE.

6. ProMED-Mail. 2018. CHRONIC WASTING DISEASE, CERVID - FINLAND: FIRST CASE, MOOSE. https://www.promedmail.org/post/5684473. Accessed

7. Benestad SL, Mitchell G, Simmons M, Ytrehus B, Vikoren T. 2016. First case of chronic wasting disease in Europe in a Norwegian free-ranging reindeer. Vet Res 47:88.

8. USGS-NWHC. Distribution of Chronic Wasting Disease in North America. https://www.usgs.gov/media/images/distribution-chronic-wasting-disease-north-america-0. Accessed 06/01/2021.

9. Sohn HJ, Kim JH, Choi KS, Nah JJ, Joo YS, Jean YH, Ahn SW, Kim OK, Kim DY, Balachandran A. 2002. A case of chronic wasting disease in an elk imported to Korea from Canada. J Vet Med Sci 64:855–8.

10. Mathiason CK, Hays SA, Powers J, Hayes-Klug J, Langenberg J, Dahmes SJ, Osborn DA, Miller KV, Warren RJ, Mason GL, Hoover EA. 2009. Infectious Prions in Pre-Clinical Deer and Transmission of Chronic Wasting Disease Solely by Environmental Exposure. PLoS ONE 4:e5916.

11. Mathiason CK, Powers JG, Dahmes SJ, Osborn DA, Miller KV, Warren RJ, Mason GL, Hays SA, Hayes-Klug J, Seelig DM, Wild MA, Wolfe LL, Spraker TR, Miller MW, Sigurdson CJ, Telling GC, Hoover EA. 2006. Infectious prions in the saliva and blood of deer with chronic wasting disease. Science 314:133–6.

12. Haley NJ, Van de Motter A, Carver S, Henderson D, Davenport K, Seelig DM, Mathiason C, Hoover E. 2013. Prion-seeding activity in cerebrospinal fluid of deer with chronic wasting disease. PLoS ONE 8:e81488.

13. Haley NJ, Seelig DM, Zabel MD, Telling GC, Hoover EA. 2009. Detection of CWD prions in urine and saliva of deer by transgenic mouse bioassay. PLoS ONE 4:e4848.

14. Henderson DM, Davenport KA, Haley NJ, Denkers ND, Mathiason CK, Hoover EA. 2015. Quantitative assessment of prion infectivity in tissues and body fluids by real-time quaking-induced conversion. J Gen Virol 96:210–9.

15. Henderson DM, Denkers ND, Hoover CE, Garbino N, Mathiason CK, Hoover EA. 2015. Longitudinal Detection of Prion Shedding in Saliva and Urine by Chronic Wasting Disease-Infected Deer by Real-Time Quaking-Induced Conversion. J Virol 89:9338–47.

16. Henderson DM, Tennant JM, Haley NJ, Denkers ND, Mathiason CK, Hoover EA. 2017. Detection of chronic wasting disease prion seeding activity in deer and elk feces by real-time quaking-induced conversion. J Gen Virol 98:1953–1962.

17. Henderson DM, Manca M, Haley NJ, Denkers ND, Nalls AV, Mathiason CK, Caughey B, Hoover EA. 2013. Rapid Antemortem Detection of CWD Prions in Deer Saliva. PLoS One 8:e74377.

18. Tamguney G, Miller MW, Wolfe LL, Sirochman TM, Glidden DV, Palmer C, Lemus A, DeArmond SJ, Prusiner SB. 2009. Asymptomatic deer excrete infectious prions in faeces. Nature 461:529–32.

19. Tamguney G, Richt JA, Hamir AN, Greenlee JJ, Miller MW, Wolfe LL, Sirochman TM, Young AJ, Glidden DV, Johnson NL, Giles K, DeArmond SJ, Prusiner SB. 2012. Salivary prions in sheep and deer. Prion 6:52–61.

20. Miller MW, Williams ES. 2003. Prion disease: horizontal prion transmission in mule deer. Nature 425:35–6.

21. Miller MW, Williams ES, Hobbs NT, Wolfe LL. 2004. Environmental sources of prion transmission in mule deer. Emerg Infect Dis 10:1003–6.

22. Mathiason CK, Hayes-Klug J, Hays SA, Powers J, Osborn DA, Dahmes SJ, Miller KV, Warren RJ, Mason GL, Telling GC, Young AJ, Hoover EA. 2010. B cells and platelets harbor prion infectivity in the blood of deer infected with chronic wasting disease. J Virol 84:5097–107.

23. Concha-Marambio L, Pritzkow S, Moda F, Tagliavini F, Ironside JW, Schulz PE, Soto C. 2016. Detection of prions in blood from patients with variant Creutzfeldt-Jakob disease. Sci Transl Med 8:370ra183.

24. Llewelyn CA, Hewitt PE, Knight RS, Amar K, Cousens S, Mackenzie J, Will RG. 2004. Possible transmission of variant Creutzfeldt-Jakob disease by blood transfusion. Lancet 363:417–21.

25. Calvente E, Pelletier S, Banfield J, Brown J, Chinnici N. 2020. Prevalence of Winter Ticks (Dermacentor albipictus) in Hunter-Harvested Wild Elk (Cervus canadensis) from Pennsylvania, USA (2017-2018). Vet Sci 7.

26. Ellingwood DD, Pekins PJ, Jones H, Musante AR. 2020. Evaluating moose *Alces alces* population response to infestation level of winter ticks *Dermacentor albipictus*. Wildlife Biology 2020.

27. McLaughlin RF, Addison EM. 1986. Tick (Dermacentor albipictus)-induced winter hair-loss in captive moose (Alces alces). J Wildl Dis 22:502–10.

28. Severud WJ, DelGiudice GD. 2016. Potential Vertical Transmission of Winter Ticks (Dermacentor albipictus) from Moose (Alces americanus) Dams to Neonates. J Wildl Dis 52:186–8.

29. Swei A, O’Connor KE, Couper LI, Thekkiniath J, Conrad PA, Padgett KA, Burns J, Yoshimizu MH, Gonzales B, Munk B, Shirkey N, Konde L, Ben Mamoun C, Lane RS, Kjemtrup A. 2019. Evidence for transmission of the zoonotic apicomplexan parasite Babesia duncani by the tick Dermacentor albipictus. Int J Parasitol 49:95–103.

30. Ewing SA, Panciera RJ, Kocan KM, Ge NL, Welsh RD, Olson RW, Barker RW, Rice LE. 1997. A winter outbreak of anaplasmosis in a nonendemic area of Oklahoma: a possible role for Dermacentor albipictus. J Vet Diagn Invest 9:206–8.

31. Haley NJ, Henderson DM, Donner R, Wyckoff S, Merrett K, Tennant J, Hoover EA, Love D, Kline E, Lehmkuhl AD, Thomsen BV. 2020. Management of chronic wasting disease in ranched elk: conclusions from a longitudinal three-year study. Prion 14:76–87.

32. Haley NJ, Henderson D, Wyckoff S, Tennant J, Hoover E, Love D, Kline E, Lehmkuhl AD, Thomsen BV. 2018. Chronic wasting disease management in ranched elk using rectal biopsy testing. Prion 12:93–108.

33. Cheng YC, Hannaoui S, John TR, Dudas S, Czub S, Gilch S. 2016. Early and Non-Invasive Detection of Chronic Wasting Disease Prions in Elk Feces by Real-Time Quaking Induced Conversion. PLoS One 11:e0166187.

34. Hoover CE, Davenport KA, Henderson DM, Denkers ND, Mathiason CK, Soto C, Zabel MD, Hoover EA. 2017. Pathways of Prion Spread during Early Chronic Wasting Disease in Deer. J Virol 91.

35. Cooper SK, Hoover CE, Henderson DM, Haley NJ, Mathiason CK, Hoover EA. 2019. Detection of CWD in cervids by RT-QuIC assay of third eyelids. PLoS One 14:e0221654.

36. Tennant JM, Li M, Henderson DM, Tyer ML, Denkers ND, Haley NJ, Mathiason CK, Hoover EA. 2020. Shedding and stability of CWD prion seeding activity in cervid feces. PLoS One 15:e0227094.

37. Haley NJ, Donner R, Henderson DM, Tennant J, Hoover EA, Manca M, Caughey B, Kondru N, Manne S, Kanthasamay A, Hannaoui S, Chang SC, Gilch S, Smiley S, Mitchell G, Lehmkuhl AD, Thomsen BV. 2020. Cross-validation of the RT-QuIC assay for the antemortem detection of chronic wasting disease in elk. Prion 14:47–55.

38. USDA. 2021. National Scrapie Eradication Program March 2021 Monthly Report. APHIShttps://www.aphis.usda.gov/animal_health/animal_diseases/scrapie/downloads/monthly_scrapie_report.pdf,

39. Authority EFS. 2020. The European Union summary report on surveillance for the presence of transmissible spongiform encephalopathies (TSE) in 2019. EFSA Journal 18:e06303.

40. Hannaoui S, Schatzl HM, Gilch S. 2017. Chronic wasting disease: Emerging prions and their potential risk. PLoS Pathog 13:e1006619.

41. Haley N, Mathiason C, Zabel MD, Telling GC, Hoover E. 2009. Detection of sub-clinical CWD infection in conventional test-negative deer long after oral exposure to urine and feces from CWD+ deer. PLoS ONE 4:e7990.

42. Kong Q, Huang S, Zou W, Vanegas D, Wang M, Wu D, Yuan J, Zheng M, Bai H, Deng H, Chen K, Jenny AL, O’Rourke K, Belay ED, Schonberger LB, Petersen RB, Sy MS, Chen SG, Gambetti P. 2005. Chronic wasting disease of elk: transmissibility to humans examined by transgenic mouse models. J Neurosci 25:7944–9.

43. LaFauci G, Carp RI, Meeker HC, Ye X, Kim JI, Natelli M, Cedeno M, Petersen RB, Kascsak R, Rubenstein R. 2006. Passage of chronic wasting disease prion into transgenic mice expressing Rocky Mountain elk (Cervus elaphus nelsoni) PrPC. J Gen Virol 87:3773–80.

44. Tamguney G, Giles K, Bouzamondo-Bernstein E, Bosque PJ, Miller MW, Safar J, DeArmond SJ, Prusiner SB. 2006. Transmission of elk and deer prions to transgenic mice. J Virol 80:9104–14.

45. Raymond GJ, Raymond LD, Meade-White KD, Hughson AG, Favara C, Gardner D, Williams ES, Miller MW, Race RE, Caughey B. 2007. Transmission and adaptation of chronic wasting disease to hamsters and transgenic mice: evidence for strains. J Virol 81:4305–14.

46. Meyerett C, Michel B, Pulford B, Spraker TR, Nichols TA, Johnson T, Kurt T, Hoover EA, Telling GC, Zabel MD. 2008. In vitro strain adaptation of CWD prions by serial protein misfolding cyclic amplification. Virology 382:267–76.

47. Kurt TD, Perrott MR, Wilusz CJ, Wilusz J, Supattapone S, Telling GC, Zabel MD, Hoover EA. 2007. Efficient in vitro amplification of chronic wasting disease PrPRES. J Virol 81:9605–8.

48. Haley NJ, Richt JA, Davenport KA, Henderson DM, Hoover EA, Manca M, Caughey B, Marthaler D, Bartz J, Gilch S. 2018. Design, implementation, and interpretation of amplification studies for prion detection. Prion 12:73–82.

49. Shikiya RA, Kincaid AE, Bartz JC, Bourret TJ. 2020. Failure To Detect Prion Infectivity in Ticks following Prion-Infected Blood Meal. mSphere 5.

50. Pekins PJ. 2020. Metabolic and Population Effects of Winter Tick Infestations on Moose: Unique Evolutionary Circumstances? Frontiers in Ecology and Evolution 8.

